# Target localization across saccades and at fixation: Nontargets both facilitate and bias responses

**DOI:** 10.1101/351833

**Authors:** Xiaoli Zhang, Julie D. Golomb

**Author notes:** Correspondence should be addressed to: Xiaoli Zhang and Julie D. Golomb.

## Abstract

The image on our retina changes every time we make an eye movement. To maintain visual stability across saccades, specifically to locate visual targets, we may use nontarget objects as “landmarks”. In the current study, we compared how the presence of nontargets affects target localization across saccades and during sustained fixation. Participants fixated a target object, which either maintained its location on the screen (sustained-fixation trials), or displaced to trigger a saccade (saccade trials). After the target disappeared, participants reported the most recent target location with a mouse click. We found that the presence of nontargets decreased response error magnitude and variability. However, this nontarget facilitation effect was not larger for saccade trials than sustained-fixation trials, indicating that nontarget facilitation might be a general effect for target localization, rather than of particular importance to saccadic stability. Additionally, participants’ responses were biased towards the nontarget locations, particularly when the nontarget-target relationships were preserved in relative coordinates across the saccade. This nontarget bias interacted with biases from other spatial references, e.g. eye movement paths, possibly in a way that emphasized non-redundant information. In summary, the presence of nontargets is one of several sources of reference that combine to influence (both facilitate and bias) target localization.

## Introduction

Starting from the retina, visual information is organized spatially, according to its retinotopic (eye-centered) location. However, this raises a critical problem as we are constantly moving our eyes, and the image received by our retina is changing accordingly, which is not optimal for world-centered (spatiotopic) cognitive tasks. Hence, there is a challenge for our visual system to distinguish real changes in the world from changes on the retina purely caused by eye movements.

It has been proposed that we are able to use information from both extra-retinal and retinal sources to achieve visual stability, for example, to localize objects accurately. Extra-retinal sources include corollary discharge or efference copy signals from saccadic eye movements, including the idea that certain visual neurons can use this information to predictively remap their receptive fields, responding to stimuli in their future receptive field locations right before a saccade [1,2]. It has been argued that remapping might be able to compensate for saccade-induced motion, or link the retinal input before and after the saccade to maintain visual stability (reviewed in [3–5]).

Another source of stability -- the focus of this project -- is retinal information: i.e., visual information in the scene. One component of retinal information is the saccade target itself; it has been proposed that the saccade target provides critical information for visual stability [6–8]. Another retinal component comes from other nontarget objects that appear in the visual field, for example a visual background [6] or frame [9], or other objects that can act as “landmarks” to influence target localization across saccades as well as at fixation [10–13].

Here we use the term “nontarget” to refer to visual objects in a display that are presented alongside a “target” object that acts as the fixation or the saccade goal. Researchers often use the terms “landmarks” or “distractors” to refer to objects presented alongside task targets that influence performance on various tasks. The term “landmark” has been mainly used in fields studying complex real-world tasks such as spatial navigation, and there is a large amount of evidence showing an important role of landmarks in performing navigation tasks (e.g., reviewed in [14]). The word “distractor” is often seen in visual attention studies, for example the influence of different types of distractor items during visual search (e.g., [15,16]). In order to avoid any confusion brought by the existing investigations of these two terms in other fields, here we use the term “nontargets”. Hypothetically these nontargets may work as “landmarks” (i.e., facilitation) or “distractors” (i.e., impairment) in target localization tasks; we use “nontargets” to remain neutral and explore both of these possibilities in our study.

Previous studies have investigated the role of nontargets in visuospatial processing in different ways. When participants were asked to saccade to a stimulus flashed during an initial eye movement, their saccade was more accurate when an egocentric cue from a visual nontarget was available [17]. It was also found that the existence of a nontarget as a visual landmark can help guide eye movements to memorized target locations more precisely, showing nontarget facilitation for the memory-based saccade execution [18]. Moreover, the presence of stable nontarget landmarks has been shown to improve detection of target displacement during fixation [19] as well as across saccades ([20] using biological-motion stimuli; [21] using bystander configuration), although nontarget landmarks have failed to facilitate visuospatial tasks in some other domains, such as intrasaccadic perception of relative motion [22].

Importantly, nontargets may influence more than just localization accuracy. For example, in target displacement detection tasks, if nontargets displace transsaccadically, it can induce illusory target displacement [10]. In this study, minor displacements of the nontargets (“landmarks” in the original paper) systematically shifted participants’ perception of target displacement, demonstrating that nontargets have an important effect on post-saccadic localization processing, presumably by acting as a stable reference point in trans-saccadic memory; in other words, any change in visual information (specifically, relative position information; also see [23]) compared with pre-saccadic memory was perceived as target displacement, regardless of whether the target actually displaced. This landmark effect may be present both during trans-saccadic tasks and at fixation [12,24].

Even when stable, nontargets can also interfere with accurate localization of targets. A phenomenon called compression of space shows that objects tend to be systematically mislocalized around the time of a saccade, such that objects are perceived to be closer to the saccade endpoint than they actually are [25], and likewise the localization of saccade targets can be compressed towards nearby nontarget objects [26]. This mislocalization might result from a “convergent remapping” component of the neuronal remapping process across saccades [27–29], although some other studies suggest that saccade might not be necessary for compression to occur [30]. This bidirectional compression indicates that the location information of nontarget objects may be integrated with target localization, even if nontarget objects only flash briefly. The idea that nontarget location information can interact with or distort target localization has also been found when nontarget objects are continuously presented along with the target. For example, Sheth & Shimojo found that during sustained fixation participants mislocalized a peripheral target as closer to a salient, unfixated bar, which acted as a visual marker [13].

In sum, the previous literature has found that the presence of visual landmarks/nontargets may help to localize targets and detect target displacement, as well as potentially bias localization and perceived target displacement. However, most studies have focused on either one effect or the other, or when they have looked at both (e.g. [13]), it has been in the context of peripheral target localization. In the current study, we focus on the localization process of the fixation/saccade target. This is because the saccade target is often critically involved in cognitive processes after saccade execution, such as memory and action [31]; hence, processing location information of the saccade target is an essential cognitive function across saccades. Our first research goal is to ask whether the facilitation and bias effects can be integrated, and how nontarget effects interact with other influences, such as fixation/saccade-related factors. For example, It has been found that localization of a peripheral target can be systematically compressed towards both a nontarget landmark and the current fixation (i.e., “foveal bias”) [13]. When the fixation point and the visual landmark were on the opposite side of the target, the total response bias was reduced, compared to when they were both on the same side of the target, suggesting that landmarks may facilitate performance by counteracting the foveal bias. Here we systematically investigate how the localization of saccade targets is influenced by nontargets, fixation-related biases, and their interaction (e.g., when they are on the same or opposite side of the target).

Second, many patterns of results mentioned above were found regardless of whether a saccade was made or planned. This brings up the question whether nontarget objects influence or facilitate target localization during saccades more than during sustained fixation, given that saccades pose unique challenges for perceptual stability [32]. The answer will tell us more about whether/how nontargets play a particularly important role in visual stability across saccades versus perception more generally. Therefore, we directly compare nontarget effects (facilitation and bias) between saccade and no-saccade trials.

Finally, when nontargets are present during a saccade target localization task, there is also the issue of reference frames: does it matter if nontargets are presented in the same absolute location across the saccade (world-centered reference frame), or should they be manipulated in relative coordinates (eye-or saccade-target-relative reference frame)? Some studies have sought to avoid this issue; for example the nontargets were simply presented on the screen at the same time as the saccade target, but were absent during the initial fixation [10]. This design (which we refer to as the “Baseline” condition in our study) focused on the role of nontargets presented at the time the saccade was triggered. But in real-world processing, nontargets rarely just appear at the time of the saccade. In the current study, we include additional conditions where nontargets are visible from the beginning of the trial (before the saccade cue). Nontargets presented before and after the saccade could remain in the same absolute location on the environment/screen (the “Absolute” condition), or remain in the same location relative to the saccade target (the “Relative” condition). Although the former case is very common and intuitive in daily experiences, many studies have suggested that the latter contains the critical information for nontargets to take effect as landmarks, at least when using displacement judgment tasks ([10]; also reviewed in [33]. It has also been found that there might be attention and/or memory benefits for relative spatial location or retinotopic coordinates across saccades, compared to absolute spatial location or spatiotopic coordinates [34–38], although other studies have found evidence for nonretinotopic processing [39–41]. However, it hasn’t been directly addressed whether stable nontargets in relative coordinates to the target would provide larger facilitation than in other reference frames.

In our project, we employed a modification of target localization tasks used in the literature, where instead of detecting trans-saccadic displacement, we simply had participants perform a target localization task by indicating target location with a mouse click (similar to [13]). Moreover, the more robust free-report task (compared to a two-alternative forced choice) allows us to measure with the response distribution not only whether target localization is facilitated or impaired under different nontarget conditions, but also whether and how much the localization reports are spatially biased by the presence of nontargets (and other factors). We tested target localization under the following conditions: Saccade presence (sustained-fixation vs saccade trials), Nontarget number (0, 1 or 2 nontargets), Congruency of the nontarget location with the initial fixation location (on the same side or opposite sides in relation to the final target) and Reference frame across saccades (Relative: the same location relative to the target; Absolute: the same absolute location on the display screen; and Baseline: not presented before the saccade target). Each reference frame condition was tested in separate experiments; within each experiment all other conditions were intermixed. We hypothesized that the presence of nontarget objects accompanying the target would both facilitate and perhaps bias target localization responses, with our main goal to investigate how this nontarget information interacts with saccade-related information, in different locations and reference frames.

## Materials and Methods

### Participants

An independent set of sixteen subjects participated in each of the three experiments (E1: 12 females, 4 males, mean age 19.06, range 18-23; E2: 9 females, 7 males, mean age 19.44, range 18-24; E3: 8 females, 8 males, mean age 20, range 18-24). All subjects reported normal or corrected-to-normal vision. They gave informed consent and were compensated with course credit or payment. The study protocols were approved by the Ohio State University Behavioral and Social Sciences Institutional Review Board.

Sample size was chosen based on a power analysis of an independent pilot experiment similar to the current study. For the main effect of nontarget (NT) number (0, 1, 2) on response error magnitude, the pilot dataset (N=16) had an effect size of *η_p_^2^*=0.493, and the power to detect such an effect was estimated as .999. We thus set N=16 as the sample size for all experiments.

### Apparatus

The experiment was run using Psychtoolbox [42] in Matlab (MathWorks). Stimuli were presented on a 21-in. CRT monitor with a refresh rate of 85 Hz. Participants were seated 61 cm in front of the monitor in a dark testing room, with a chinrest for eye-tracking purposes.

### Eye-tracking

Eye positions were recorded throughout the experiment using an Eyelink 1000 Eye Tracker at 500 Hz. Eye position data were used to ensure the participants kept their eyes on the target, and to measure saccade trajectories and latencies. If they were not fixating at the correct location, a “Fixation Error!” message was shown on the screen, the current trial failed immediately, and the next trial started. The failed trials were re-run in a random order later in the block. Saccades were identified and analyzed using custom Matlab code as described below.

### Task procedure

Three experiments were run to look at the effect of nontargets on target localization across saccades and at fixation. The paradigm is shown in Fig 1.

**Fig 1.**
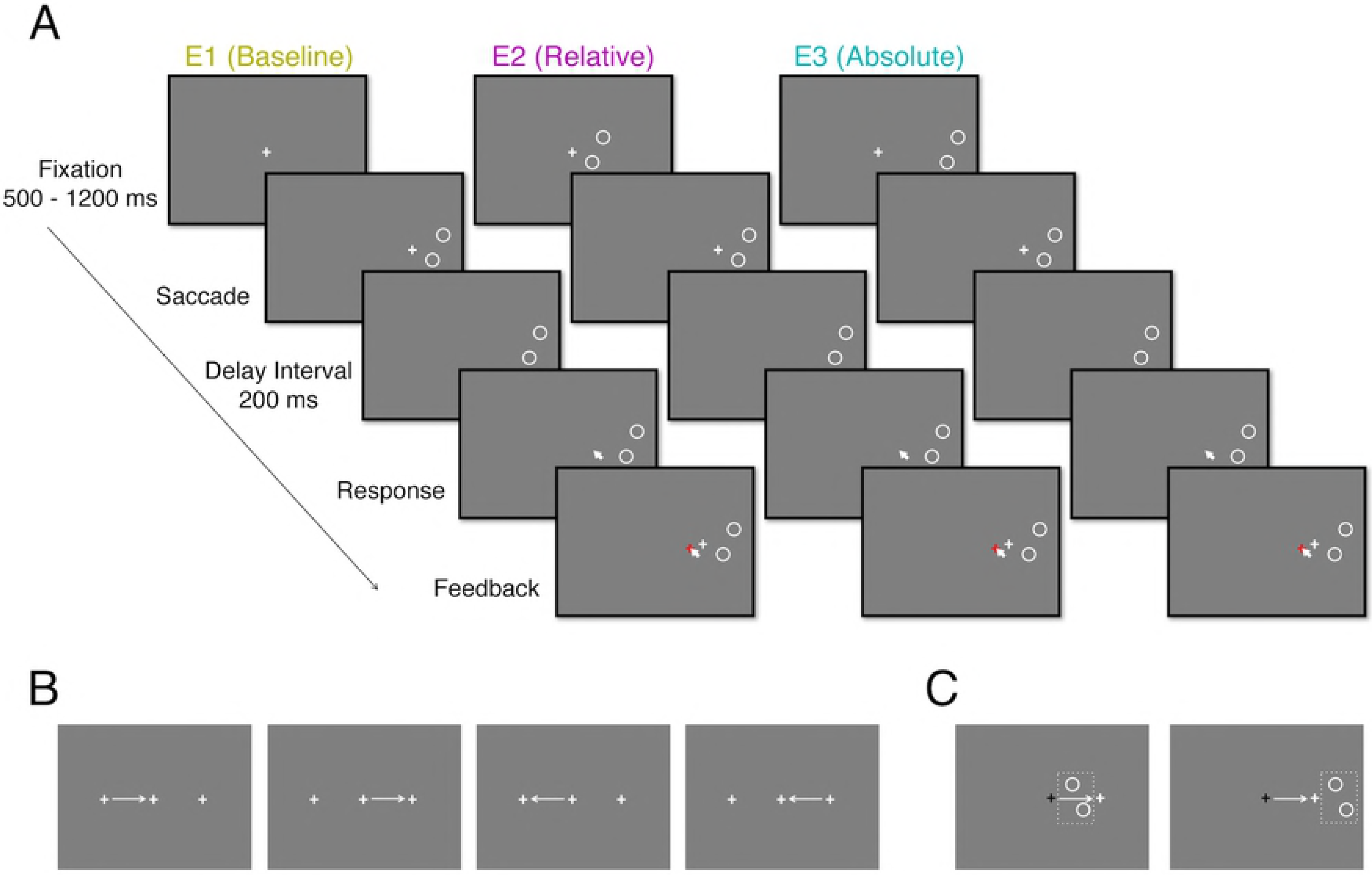
Experiment Paradigm. A) Sample trial sequence for each of the three experiments. Each example shows a rightward saccade trial with 2 nontargets (white circles) on the right side. In E1 the nontargets appear at the same time as the saccade cue. In E2 and E3 the nontargets are present from the beginning of the trial and maintain the same locations relative to the saccade target (E2) or absolute screen location (E3). After subjects successfully complete the saccade, the target is removed for a 200ms delay, and then participants make a response by moving the cursor and clicking at the remembered final target location. Feedback (a red cross at the reported location and a white cross at the correct location) was presented after response. B) Schematic indicating the different possible saccade patterns. Crosses indicate the three possible target locations, and arrows show saccade patterns; note that the actual target locations were jittered on each trial. C) Schematic showing different conditions of congruency on a sample saccade trial. Black crosses indicate the initial fixation and white crosses indicate the saccade target location. For a rightward saccade, NTs could appear either to the left or right of the final target. Left panel: when NTs and the initial fixation location were to the left of the final target location, this would be a same-side condition. Right panel: when NTs were to the right and the initial fixation location was to the left of the final target location, this would be an opposite-side condition. Dashed rectangle indicates the possible extent of the NT region; the actual nontargets (circles) were randomly presented in that rectangle region in each trial.

For all experiments, participants began each trial by fixating a white cross sized 0. 2°×0.2° (the target) on a constant gray background, RGB (127, 127, 127). The horizontal location of the target was randomized among three possible locations - 4° left of, 4° right of, and on the vertical midline, with 0° - 0.25° additional random jitter. The vertical location was also jittered within 0.25° above or below the horizontal midline of the screen. Once participants were fixating (i.e., the eye location stayed within 1.5° range of the target), the target stayed visible for a variable period of 500 to 1000 ms. On saccade trials (50% of all trials), the target then jumped to an adjacent location to trigger a horizontal saccade of 4° (Fig 1B). The saccade end time was determined when the participants’ eye position was within 1.5° range from the saccade target and the velocity of the eye movement was below 30°/s [43]. Trials failed immediately if the saccade was not completed within 3 s after the target jump.

After the saccade was detected as completed, the target was removed for 200 ms. This means that the target was removed post-saccadically, but not midflight. Note that the goal of this design is to not to investigate trans-saccadic perception per se, but how target localization before and after saccades is affected by the presence of nontargets. On no-saccade trials, the target was removed from its initial location after a delay analogous to saccade latency (250 - 300 ms). Following this 200ms blank interval, a beep sound occurred to instruct participants to respond by moving the cursor to the remembered target location - the center of the cross. The cursor was presented on the screen at a random starting point 0.5° to 1° away from the target, to eliminate the effect of cursor location across trials. Participants clicked the left button to register their response. Feedback with the correct and reported location was shown for 1000 ms.

On some trials, nontarget objects (white empty circles of 0.2° radius) were also presented during the trial: trials were equally distributed among 0, 1, or 2 NTs. Participants were told that they should complete the task on the target cross, and that the circles were irrelevant to their task. In Experiment 1 (Baseline), nontargets appeared on the screen simultaneously with the saccade target (second fixation cue), or after an analogous delay on no-saccade trials. In Experiments 2 & 3, nontargets appeared at the beginning of the trial, and remained on the screen throughout the trial in either “Relative” (Experiment 2) or “Absolute” (Experiment 3) reference frames. In Experiment 2, nontargets remained in the same location relative to the fixation cross (i.e., they moved with the saccade target; see Fig 1). In Experiment 3, nontargets remained in the same absolute location on the screen across the saccade.

In all three experiments, we designed the *NT location* conditions to be either to the left or right of the target’s final position, and thus either on the same side or opposite side as the initial fixation on saccade trials (Fig 1C). The actual NT locations were randomized for each trial within an imaginary vertical rectangle zone of 1°× 2°, centered 2° to the left or right of the target. This means that on trials with 2 NTs, these two NTs were both presented on the same side of the target. In the Baseline experiment, NTs were presented when the target appeared in its final position, centered 2° to the left or right of that final target location. In the Relative experiment, the NTs first appeared centered 2° to the left or right of the initial target location, and moved with the target to remain in the same relative location. Note that because the NTs moved with the target instead of the eyes, we call this condition “relative” instead of “retinotopic”. In the Absolute experiment, we included three different scenarios (S3A Fig). For rightward saccades, these scenarios were as follows: (a) the NTs appeared centered 2° to the right of the initial target position, which made them 2° to the left of the final target position (“near-near”); (b) the NTs appeared 2° to the left of the initial target position, meaning 6° to the left of the final target position (“near-far”); (c) the NTs appeared 6° to the right of the initial target position, meaning 2° to the right of the final target position (“far-near”). It is an intrinsic confound in the Absolute experiment that the distance between NTs and the target could not be kept at 2° before and after a saccade and still include a mix of same-side and opposite-side conditions. Therefore, we included all three distance conditions described above to cover both same-side and opposite-side conditions in the Absolute experiment. For the main analyses, we collapsed across these three distance conditions. Separate results for the three distance conditions are shown in the supplementary materials.

For all experiments, participants completed a practice block, and then there were 12 main task blocks, 48 trials each. These 48 trials were equally distributed among the 2 saccade presence (no-saccade and saccade) × 3 NT number (0, 1 and 2 NTs) × 2 NT location (same and opposite side relative to initial fixation). A minimum of 8 blocks was set as a threshold for the data to be included in analyses (some participants could not complete the full 12 blocks in the allotted 1.5-hour session due to eye tracking difficulty). Each subject thus completed 32-48 trials per critical condition described above.

### Data processing and analyses

Data were processed with custom Matlab (version 2015b) code and analyzed in JASP [44]. Trials with unreasonably long reaction time (>7s) or unreasonably large localization error (>1.5°) were discarded. The latter means that the situation where participants mistook the NT location as the target location was excluded. The discarded trials took up less than 0.2% of all trials in each experiment.

The conditions we analyzed included saccade presence (no-saccade and saccade), NT number (0, 1 and 2 NTs), and NT location (same and opposite side relative to initial fixation location). Each of these conditions was tested within each experiment (within-subjects), and compared across experiments (between-subjects), which varied reference frame.

Our primary goal was to assess how the above factors influence target localization performance; thus, the analyses primarily focus on the participants’ mouse responses (though we include some additional analyses of eye-tracking data in the supplementary materials). We first investigated how making saccades influences target localization by comparing saccade versus no-saccade trials; then how NTs influence target localization by comparing trials with zero, one and two NTs; and finally, if/how these saccade and NT influences interact by analyzing saccade trials with NTs. We used three measurements to quantify target localization outcomes: 1) how accurate participants’ responses were, by calculating the mean error magnitude as the distance (i.e., absolute value) between the reported and correct target location; 2) how variable participants’ responses were, by calculating the root mean squared distance (RMSD) for each condition of interest for each subject; 3) how biased participants’ responses were, by calculating the mean directional error vector along the horizontal axis along which saccades and NT locations were manipulated.

Specifically, RMSD was calculated using the formula:

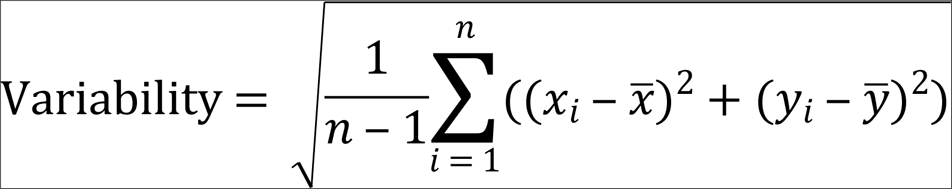

where for each subject each condition, (*x*_*i*_, *y*_*i*_) is the response coordinates for trial *i*, centered around the actual target location; (*x̅*, *y̅*) is the average coordinates of all responses in that condition; *n* is the number of trials, and the denominator (*n*-1) is the degree of freedom to get an unbiased estimate.

All of the above three measurements were calculated in units of visual angle. We used ANOVAs and t-tests for statistical analyses; effect sizes were calculated using *η*_*p*_^*2*^ and Cohen’s *d*. Greenhouse-Geisser correction for violations of sphericity and Holm-Bonferroni correction for multiple comparisons were used when necessary.

## Results

Our research question focused on how saccades and nontargets influence target localization independently and interactively.

A descriptive plot of participants’ responses is depicted in Fig 2, where a scatter plot of participants’ responses in each trial is plotted relative to the correct target location and saccade / NT directions, and 95% confidence ellipses of response error summarize the accuracy, precision, and bias of these responses (error ellipses calculated according to [45]). Statistical comparisons for each question of interest follow in the sections below.

**Fig 2.**
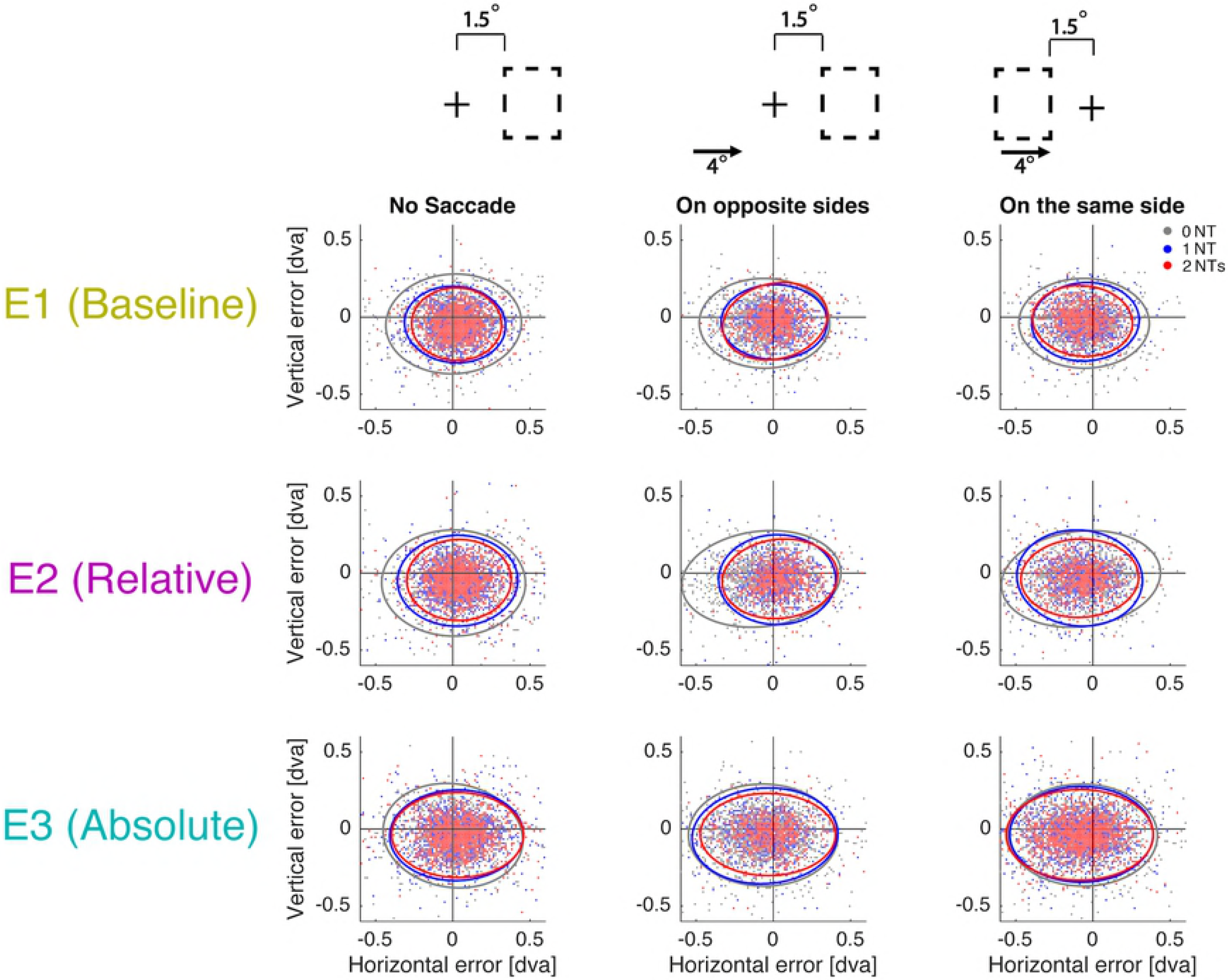
Scatter plots of participants’ localization errors across conditions in each experiment. X and y axes represent response error (in degree visual angle, dva) on horizontal and vertical axes respectively. (0,0) is the correct target location. Error ellipses show the 95% confidence interval of covariance between response errors on x and y axes. All trials were aligned according to the schematics above each column. The cross indicates the final target; the dashed rectangle indicates the range of nontarget (NT) location; the arrow indicates saccade direction. Note that the schematics are not drawn to scale or in the same scale as the scatter plots; for reference, the majority of responses were made within 0.5deg of the target, the closest NTs were 1.5deg from the target, and the initial fixation was 4deg from the target. The first column shows no-saccade trials. The second and third columns show saccade trials when NTs and the initial fixation location were on opposite sides of the target and when they were on the same side of the target, respectively. Rows correspond to the 3 experiments. Within each plot, data are shown for 0, 1, and 2 NTs, including all participants for visualization. N=16 for each experiment.

### Accuracy of target localization

We first looked at the effects of saccades and NTs on overall target localization accuracy, measured by the mean magnitude of error (distance) between the correct and reported locations. Note that this initial measure doesn’t include information on which direction the participants made the error. Data were submitted to a 2 (saccade presence: 0, 1) × 3 (NT number: 0, 1, 2) × 3 (experiment: 1, 2, 3) mixed-design ANOVA.

The results showed a significant main effect of saccade presence, *F*(1,45)=15.351, p<.001, *η*_*p*_^*2*^=.254, indicating that the error magnitude was larger in saccade trials than no-saccade trials. There was also a main effect of NT number, *F*(1.503,67.662)=46.809, *p*<.001, *η*_*p*_^*2*^=.510, that increasing the number of NTs decreased the error magnitude. There was no significant interaction between saccade presence and NT number, *F*(1.647,74.111)=0.059, *p*=.913, *η*_*p*_^*2*^=.001, indicating that the influence of NTs on target localization accuracy was similar for both saccade and no-saccade trials.

Do these influences of NTs and saccades vary across our different experiments? In Experiment 1 (baseline), NTs were presented at the same time as the saccade target, whereas in Experiments 2 and 3 NTs were presented before the saccade target, in relative (same location relative to target) and absolute (same absolute location on screen) coordinates, respectively. We found a significant interaction between experiment and NT number, *F*(3.005,67.622)=4.201, *p*=.009, *η*_*p*_^*2*^=.157, but no significant main effect of experiment nor interaction between saccade presence and experiment, *F*(2,45)=1.338, *p*=.273, *η*_*p*_^*2*^=.056, *F*(2,45)=1.211, *p*=.307, *η*_*p*_^*2*^=.051. There was no significant three-way interaction between saccade presence, NT number and experiment, *F*(3.294,74.111)=1.833, *p*=.143, *η*_*p*_^*2*^=.075. Fig 3A illustrates the NT number × experiment interaction. The presence of NTs decreased error in all experiments, but this NT facilitation effect was greater for the baseline and relative conditions (E1 and E2) compared to the absolute condition (E3). Using the zero NT trials as a baseline for each experiment, we calculated the “NT facilitation” effect for NT1 and NT2 trials for each of the 3 experiments. A 2 (NT number: 1, 2) × 2 (saccade presence: 0, 1) × 3 (experiment: 1, 2, 3) mixed-design ANOVA found a significant main effect of NT number, *F*(1,45)=6.914, *p*=.012, *η*_*p*_^*2*^=.133, showing greater facilitation with two nontargets than one nontarget, along with a main effect of experiment, *F*(2,45)=5.206, *p*=.009, *η*_*p*_^*2*^=.188. Post hoc t-tests between experiments showed that NT facilitation was not significantly different between baseline and relative conditions, *t*(30)=-0.447, *p*=.658, Cohen’s *d*=-0.158, but that in both baseline and relative conditions facilitation effects were significantly larger than in the absolute condition (*t*(30)=-3.920, *p*<.001, Cohen’s *d*=-1.386 and *t*(30)=-2.477, *p*=0.019, Cohen’s *d*=-0.876, respectively). It is possible that some of these experiment effects could be driven by distance effects - i.e. in the absolute condition some nontargets were located further from the target (see methods). We then restricted Absolute trials to the subset that matched the distance of relative NTs (i.e., “near-near” condition), and we still found a significant difference between Absolute and Relative facilitation, *F*(1,15)=6.712, *p*=.020, *η*_*p*_^*2*^=.309 (additional results in the supplementary materials).

**Fig 3.**
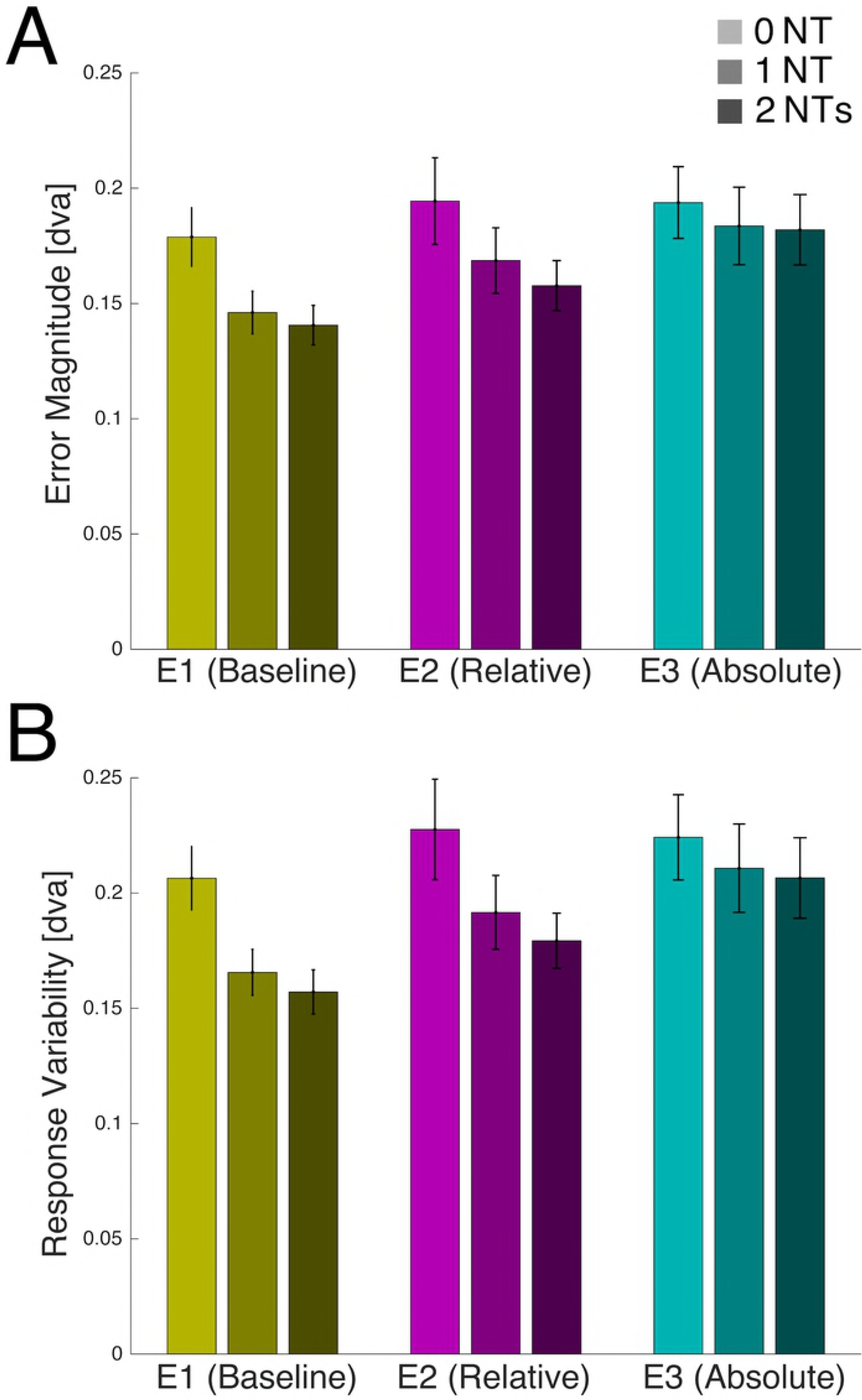
Target localization performance. Target localization error magnitude (A) and response variability (B) as a function of NT number, in each of the three experiments. Data are collapsed across saccade and no-saccade trials. N=16 for each experiment. Error bars are SEM.

### Variability of responses

We next examined another important measurement of target localization performance, the variability of the responses, quantified using RMSD.

We did similar analyses as above, using a 2 (saccade presence: 0, 1) × 3 (NT number: 0, 1, 2) × 3 (experiment: 1, 2, 3) mixed-design ANOVA, and found similar patterns. There was a significant main effect of NT number, *F*(1.625,73.108)=52.783, *p*<.001, *η*_*p*_^*2*^= 540, where NTs reduced response variability. Making a saccade significantly increased response variability, *F*(1,45)=13.133, *p*<.001, *η*_*p*_^*2*^=.226. There was no significant interaction between saccade presence and NT number, *F*(1.670,75.132)=2.059, *p*=.142, *η*_*p*_^*2*^=.044.

There was no significant interaction between saccade presence and experiment, *F*(2,45)=0.955, *p*=.392, *η*_*p*_^*2*^=.041. The NT number × experiment interaction was significant, *F*(3.249,73.108)=3.984, *p*=.009, *η*_*p*_^*2*^=.150. As shown in Fig 3B, NT facilitation affected variability in a similar way as overall accuracy. NT facilitation was present in all three experiments, but was greater for the baseline and relative conditions (E1 and E2) compared to the absolute condition (E3), *F*(2,45)=5.503, *p*=.007, *η*_*p*_^*2*^=.197, and was greater for 2NT than 1NT, *F*(1,45)=7.300, *p*#.010, *η*_*p*_^*2*^=.140. Similar to accuracy analyses above, when restricting to trials in which NT distance was comparable across experiments, Relative facilitation was still greater than Absolute facilitation, *F*(1,15)=7.405, *p*#.016, *η*_*p*_^*2*^=.331.

### Spatial response biases

So far, we assessed the performance of target localization in terms of error magnitude and response variability, and found that the presence of nontargets decreased both measurements; i.e. nontargets improved target localization performance on both saccade and no-saccade trials. However, it should be noted that these two measurements ignored the directional information of participants’ responses. That is, were errors randomly distributed around the correct location, or was there systematic variability? There could be two ways in which directional error might be informative here: First, there might be a difference in horizontal versus vertical error magnitudes (particularly because in our paradigm, saccade direction was only manipulated along the horizontal axis). Second, we can ask whether the saccade direction and/or location of the NTs on a given trial might systematically *bias* the reported target location, e.g. toward or away from the NTs or initial fixation.

To address the first question, we performed the same analysis as above for mean error magnitude, but now separately for horizontal and vertical error magnitude. The increase in error on saccade versus no-saccade trials happened only along the horizontal axis; interestingly, making a saccade actually decreased the error along vertical axis (horizontal: *F*(1,45)=28.288, *p*<.001, *η_p_^2^*=.386; vertical: *F*(1,45)=10.791, *p*#.002, *η*_*p*_^*2*^=.193). NT facilitation happened along both horizontal and vertical axes. However, the experiment × NT interaction was only found along the horizontal axis (horizontal: *F*(3.017,67.893)=5.009, *p*#.003, *η*_*p*_^*2*^=.182; vertical: *F*(3.592,80.825)=0.909, *p*#.454, *η_p_^2^*=.039). Similar patterns were found for response variability: making a saccade increased response variability only along the horizontal axis (horizontal: *F*(1,45)=18.362, *p*<.001, *η*_*p*_^*2*^=.290; vertical: *F*(1,45)=0.740, *p*#.394, *η*_*p*_^*2*^=.016); and NT facilitation existed along both horizontal and vertical axes, but interacted with experiment only along the horizontal axis (horizontal: *F*(3.279,73.781)=5.065, *p*#.002, *η*_*p*_^*2*^=184; vertical: *F*(3.782,85.098)=0.542, *p*#.695, *η*_*p*_^*2*^=.024).

Because saccades were only executed along the horizontal axis, and the NT x experiment interaction was also specific to the horizontal axis, for our second question (i.e., spatial bias), we focused primarily on horizontal directional error. To enable us to look at the joint influence of saccade and NT biases, we simplified the location of NTs into whether they were presented in the same horizontal direction as the initial fixation (Same) or opposite horizontal direction (Opposite).

#### Does saccade direction bias target localization?

To isolate a potential saccade-related bias, we first restricted our analyses to trials with zero NTs (Fig 4B and 4C, when NT number is zero in saccade trials; also S1B Fig). We aligned each trial’s data so that a positive error vector would mean bias towards the initial fixation location on saccade trials (and towards right on no-saccade trials). A 2 (saccade presence: 0, 1) × 3 (experiment: 1, 2, 3) mixed-design ANOVA found a significant main effect of making a saccade, *F*(1,45)=54.863, *p*<.001, *η*_*p*_^*2*^=.549, with participants’ responses more biased on saccade than no-saccade trials. Post-hoc tests revealed that on saccade trials, target localization (mouse) responses were significantly biased towards the initial fixation location (compared to zero bias: *t*(47)=-7.482, *p*<.001, Cohen’s *d*#-1.080), while the bias on no-saccade trials was not significantly different from zero *t*(47)=0.879, *p*#.384, Cohen’s *d*#0.127. There was no significant main effect of or interaction with experiment, *F*(2,45)=0.311, *p*#.734, *η*_*p*_^*2*^=.014, *F*(2,45)=0.351, *p*#.706, *η*_*p*_^*2*^=.015, respectively. A supplementary analysis (S1 Fig) revealed that there was also a similar bias in saccade landing point, with the majority of saccade trials undershooting the target. However - critically - target localization (mouse response) was biased towards the initial fixation location regardless of actual saccade endpoint. We compared saccade undershoot and overshoot trials separately and found that for both saccade undershoot and overshoot trials, there was a significant localization bias in the direction of initial fixation in all experiments, *t*’s≥2.802, *p*’s≤.013, Cohen’s *d*’s≥0.700; i.e., saccade endpoint (undershoot or overshoot) impacted the magnitude of this bias, *F*(1,45)=9.102, *p*#.004, *η*_*p*_^*2*^=.168, but did not drive the effect.

**Fig 4.**
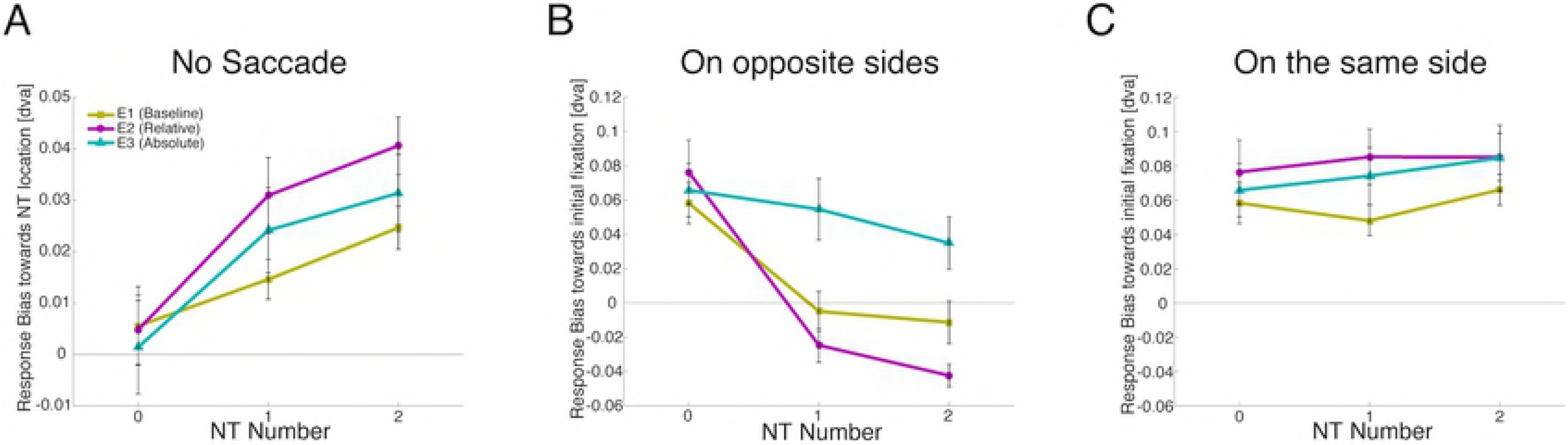
Spatial response biases. A) Response biases on no-saccade trials (NT-related bias). Positive values mean biasing towards NT location (and towards right when NT number is zero). Bias increases with NT number. B) Response biases on saccade trials when NT location and initial fixation are on the opposite sides of the target. Positive values mean biasing towards the initial fixation location. When the NT number is zero, the positive values in three experiments indicate a saccade-related response bias towards the initial fixation. NTs on the opposite side counteract this bias. C) Response biases on saccade trials when NT location and initial fixation are on the same side of the target. As in B, when the NT number is zero, the positive values in three experiments indicate a saccade-related response bias towards the initial fixation. However, NTs on the same side add little to this bias. N=16 for each experiment. Error bars are SEM.

#### Do nontargets bias target localization?

Next, to explore the potential bias from nontargets alone, we looked at no-saccade trials, comparing zero, one or two NTs (Fig 4A). We aligned the data so that a positive error vector would mean bias towards the NTs (along horizontal axis). A 3 (NT number: 0, 1, 2) × 3 (experiment: 1, 2, 3) mixed-design ANOVA found a significant main effect of NT number, *F*(1.425,64.144)=13.062, *p*<.001, *η*_*p*_^*2*^=.225. On trials where NTs were present, participants’ responses were biased towards the NT location; the bias was significant for both 1NT and 2NT, *t*(47)=5.879, *p*<.001, Cohen’s *d*#0.849, *t*(47)=9.242, *p*<.001, Cohen’s *d*#1.334, respectively, and two NTs yielded a significantly larger bias than one, *t*(47)=2.645, *p*#.011, Cohen’s *d*#0.382. There was no significant main effect of or interaction with experiment, *F*(2,45)=1.791, *p*#.179, *η*_*p*_^*2*^=.074, *F*(2.851,64.144)=0.541, *p*#.647, *η*_*p*_^*2*^=.023, respectively.

#### Joint influence of saccade and NTs

Now the key question is: how do these two sources of bias *interact* when both are present? E.g., if the biases induced by the saccade and the NTs are in the same direction, do they add together to result in a larger bias? If the sources of bias are in opposite directions, do they counteract each other? In other words, can the presence of non-targets compensate for the bias induced by the saccade? For this analysis, we separated saccade trials into cases where the initial fixation and the NTs were on opposite sides of the target (*Opposite side* condition, Fig 4B), or on the same side of the target (*Same side* condition, Fig 4C). We conducted a 3 (NT number: 0, 1, 2) × 2 (Congruency: same side, opposite side) × 3 (experiment: 1, 2, 3) mixed-design ANOVA on the saccade trials; in order to make the ANOVA feasible, we dummy-coded saccade trials with zero nontargets to be randomly assigned to the same or opposite side.

We found a significant main effect of NT number, *F*(1.375,61.869)=24.911, *p*<.001, *η*_*p*_^*2*^=.356, showing that overall the presence of nontargets biased responses towards the NT location, as before for the no-saccade trials. There was a significant main effect of congruency, *F*(1,45)=49.892, *p*<.001, *η*_*p*_^*2*^=.526, and a congruency × NT number interaction, *F*(1.593,71.665)=39.222, *p*<.001, *η*_*p*_^*2*^=.466. There were also significant Experiment × NT number and Experiment × Congruency interactions, *F*(2.750,61.869)=5.740, *p*#.002, *η*_*p*_^*2*^=.203 and *F*(2,45)=7.774, *p*#.001, *η*_*p*_^*2*^=.257, respectively. The 3-way interaction between NT number, experiment and congruency was not significant, *F*(3.185,71.665)=1.970, *p*#.123, *η*_*p*_^*2*^=.080.

To better explore these interactions, we separated the same side and opposite side trials and did a 3 (NT number: 0, 1, 2) × 3 (experiment: 1, 2, 3) mixed-design ANOVA on each. When NTs were on the same side as the initial fixation (Fig 4C), there was a relatively stable positive response bias (i.e., toward the initial fixation); there was no significant main effect of NT number or experiment, nor NT number × experiment interaction, all *F*’s≤1.905, *p*’s≥167, *η_p_^2^*’s≤.061. This implies that when NTs were presented on the same side of the target as the initial fixation, there was no additivity of the biases; the magnitude of the bias on these trials was the same as the saccade-related bias alone on 0-NT trials.

However, when NTs were on the opposite side of the target as the initial fixation (Fig 4B), we found a significant main effect of NT number, *F*(1.498,67.408)=53.383, *p*<.001, *η*_*p*_^*2*^=.543, a significant main effect of experiment, *F*(2,45)=6.180, *p*#.004, *η*_*p*_^*2*^=.215, and a significant interaction, *F*(2.996,67.408)=5.495, *p*#.002, *η*_*p*_^*2*^=.196. The addition of the NTs here seemed to counteract the saccade-related bias coming from the opposite direction, with the influence of 2 NTs significantly greater than 1 NT, *t*(47)=3.027, *p*#.004, Cohen’s *d*#0.437.

Interestingly, the degree to which the NTs counteracted the saccade-related bias varied by experiment. In the Baseline experiment (E1), the saccade-related bias appeared to be completely counteracted by the opposite-side NTs; the response bias when NTs were present was not significantly different from zero, *t*(15)=-0.713, *p*#.487, Cohen’s *d*#0.178 (post-hoc t-test collapsing across 1 and 2 NTs), suggesting equal and opposite contributions from the NT-related and saccade-related biases. In the Relative experiment (E2), the NT influence seemed to exceed the saccade-related bias; here the response bias was significantly negative (towards NTs, away from initial fixation), *t*(15)=-4.312, *p*#.002, Cohen’s *d*#1.078, in such a way that the NT-related bias overcompensated saccade-related bias. In contrast, in the Absolute experiment (E3), the NT-related bias did not fully counteract the saccade-related bias; here the response bias was still significantly positive (towards initial fixation), *t*(15)=2.809, *p*#.026, Cohen’s *d*#-0.702. For these three t-tests, P values were corrected for multiple comparisons using Holm-Bonferroni correction. This pattern of results implies that the bias induced by the presence of NTs was more influential when NTs were presented in the relative reference frame than absolute reference frame across saccades.

## Discussion

In the current study, we tested how the presence of nontargets influences target localization across saccades and during sustained fixation. Unsurprisingly, we found that target localization performance was generally worse on saccade than no-saccade trials (in terms of mean error magnitude and response variability), and the presence of nontargets improved target localization performance. The presence of nontargets exerted comparable facilitation effects on saccade trials and no-saccade trials, suggesting that the facilitation effect is a more general visual effect rather than of particular importance to saccadic stability. We also measured response bias (directional error), finding that participants’ responses were biased towards both the initial fixation location (saccade-related bias) and the NT locations. These two sources of bias interacted in an interesting way: When both sources fell on the same side of the target they were not additive, but when they fell on opposite sides of the target, the NT bias counteracted the saccade-related bias. For both facilitation and bias effects, the influence of nontargets was stronger when there were 2 NTs than 1 NT, and was weaker in the absolute than relative and baseline experiments. Below we discuss the implications of each of these findings.

### Saccade influence on target localization

A large literature has focused on the challenge of maintaining visual stability while moving the eyes around, particularly in terms of target localization abilities. In all three experiments, we found that saccades impaired performance by increasing error magnitude as well as response variability, even though the target was fixated within the fovea, where visual acuity and overt attention is the highest. The saccade-related increase in error magnitude and response variability happened only along the horizontal axis, such that the location errors become elongated along the saccade axis. This basic finding is intuitive, and is consistent with previous findings [10,13,24].

In addition to a generic saccade-related decrease in performance, we also found a systematic saccade-related bias: participants’ responses were on average biased in the opposite direction of the saccade. There are three possible sources of this saccade-related bias: bias towards the screen center, bias towards the actual saccade landing position, and/or bias towards the initial fixation location. In our design, the potential effect of screen center location was controlled - a left/right saccade could be from center to periphery on the screen or vice versa (Fig 1) - so the screen center is not likely to be the source of this saccade-related bias. The second and third possibilities, however, could both have predicted a systematic response bias in the same direction as we found: as reported above, both the eye landing position and the mouse responses were biased towards initial fixation on average. However, the analysis differentiating the influence of saccade landing position and initial fixation location revealed that while saccade landing position did modulate the magnitude of response bias, there was still a significant bias towards the initial fixation location even on overshoot trials when the actual eye position was in the opposite direction of the target. Thus, while actual current eye position may induce some bias (similar to the influence of saccade landing site on perception of the target displacement, shown in [46]), the primary source of the saccade-related response bias here seems to be the initial fixation location. Participants may have been using the pre-saccadic fixation location as a visual or oculomotor reference, and target localization responses were biased towards this reference; however, participants were not simply clicking on the location that they looked at.

Our result is consistent with a number of previous studies demonstrating a response bias towards the current and/or initial fixation locations [13,36,47]. Sheth and Shimojo found that visual memory of peripheral spatial locations can be biased towards the current fixation (i.e., “foveal bias”) over time, independent of saccade preparation or saccade execution. They proposed that this bias likely happens during encoding period when the eccentricity of the target might be underestimated [13]. A response bias towards the initial fixation location has also been found across saccades, when participants retained spatial memory of a peripheral target [36]. It should be noted that our design differed from these previous studies in that instead of a peripheral target, our target was the saccade target to be fixated on. However, we propose that the saccade-related bias in our result likely happened in a similar way as the studies mentioned above. When the saccade target location was presented on the screen while participants were still fixating on the initial fixation, the saccade target was indeed in the periphery at that time point. Due to the underestimated eccentricity during the encoding process, a biased representation of space was likely created and maintained across the saccade. Therefore, we still found “foveal bias” - bias towards the initial fixation, after the saccade was completed. Indeed, the magnitude of saccade-related bias we found (0.05°) is much smaller than the foveal bias in [13] (about 1°), and this is likely due to the acuity difference between processing foveal and peripheral targets.

### Nontarget facilitation on both error magnitude and response variability

The influence of nontargets on target localization has been investigated in many studies, including the presence of nontargets on saccade execution accuracy [18] and the effect of NT displacement on target displacement perception [10,19–21,24]. In our study, we focused on the influence of nontargets on target localization in a more systematic manner: investigating the number, location and reference frame of nontargets. We found that the presence of stable nontargets in general facilitated performance, by decreasing the mean error magnitude as well as response variability. The magnitude of NT facilitation was small in absolute terms (about 0.025° or 1 pixel), but reflected an improvement of approximately 14% of the baseline for absolute error measurement, and 12% for response variability measurement. The correct target location landed in the fovea, and there were other potential references such as the display boundaries; therefore, even an improvement of 1 pixel is a meaningful benefit provided by the presence of nontargets.

Did the NT facilitation stem from a direct effect - i.e. a more precise representation of target location - or it is possible that nontargets instead helped sustain fixation at or execute saccades to the target more accurately, which as a result could indirectly make the behavioral responses more accurate? To test this latter possibility, we analyzed the influence of nontargets on eye position accuracy (error distance between the target position and actual eye position) as well as eye position variability (RMSD of actual eye position) at the time point when the target was removed from the screen before the localization response (S2 Fig). If anything, the presence of nontargets actually increased eye position error magnitude and variability, suggesting that nontargets indeed facilitated the representation of target location.

Our results reflect the idea that nontargets perform as anchors or landmarks, so that the target localization could be done with them as relative references in space, consistent with previous literature (e.g., [10]; see later discussion on the effect of reference frame). Note that in our experiments, we did not explicitly instruct participants to use nontargets, which means that nontarget information might be processed and used by default, instead of only triggered by instruction. Our results showed that two nontargets facilitated slightly more than one, but the second nontarget did not double the facilitation. A possible reason is that in our design, the two nontargets always appeared inside one rectangle region: they were always on the same side of the target, and their distance to the target was similar (within 1.5° and 2.5° to the target location on the horizontal axis). Thus, the two NT objects might have been grouped together as a single landmark, or simply provided similar information, and therefore, the second nontarget might not have provided much additional reference beyond the first one. We also found that when the initial fixation location and nontargets were on the same side of the target, the presence of nontargets did not add on to the response bias (discussed below in more detail). This result supports a similar interpretation, that multiple sources of reference located on the same side might provide some redundant information which is relatively less useful for localization.

### No additional nontarget facilitation on saccade trials

Though nontargets facilitated target localization on both no-saccade and saccade trials, we did not find larger magnitude of NT facilitation on saccade trials compared to no-saccade trials. This means that nontargets did not provide *additional* facilitation across a saccade compared to sustained fixation, consistent across all three experiments. In the visual stability literature, landmarks are often highlighted for their role aiding stability across saccades. However, what is often less emphasized is that these NT effects may occur independently of the saccades. Yet our study is certainly not the first to report this. Deubel and his colleagues showed that a displacement of NT objects following a blank period after the saccade might lead participants to misjudge the target location. When there was no saccade, the displacement of the nontargets after the blank had a similar effect compared to saccade trials, even though during continuous presentation participants could detect target displacement without error [24]. This result pattern was replicated in [12].

What does this mean for visual stability? Based on our results as well as previous studies, we propose that nontargets may be useful references during saccades, but the effect of nontargets seems to be more general; i.e., even though saccades pose particular challenges for visual stability, nontargets may not be more helpful in saccade cases than sustained fixation.

### Bias induced by nontarget location

In addition to the nontarget facilitation effect, one of the more interesting influences of nontargets in our study was the biasing of target responses towards the nontarget locations, as well as how this bias interacts with the saccade-related response bias.

Response biases between fixation/saccade target and nontarget objects have been shown in previous studies, for example with perisaccadic compression of space [25,26,32] and other types of landmark-related bias [13]. The former paradigm used nontargets that briefly flashed around the time of a saccade, and the latter study tested target localization in the periphery, while our study tested stable nontargets and foveal target localization. We found a similar response bias towards nontarget location as the previous studies, although the magnitude of our nontarget bias was smaller compared with Sheth & Shimojo’s result in [13]. This is again likely due to more accurate visual processing in the fovea compared to the periphery.

What happened on saccade trials where the saccade-related bias and NT-related bias could both take place? When the nontarget location and the initial fixation were on opposite sides of the target, the nontarget bias combined with (i.e., counteracted) the saccade-related bias. However, we found that when the nontargets and initial fixation were on the same side, the two sources of biases did not appear to combine; in fact, the response bias was not any larger than the saccade-related bias alone (i.e., saccade trials with zero nontargets).

This result pattern we found was partially shown in Sheth and Shimojo’s study. They found that when a salient landmark was displayed on the opposite side of the fixation to the target, the response bias was reduced compared to on the same side, meaning that the landmark biased responses in the opposite direction and counteracted the foveal bias to the fixation [13]. However, in their study when the landmark-related bias and fixation-induced bias were in the same direction, the landmark was at least 42° away from the target, which likely did not induce landmark-related bias at all. Our study carefully controlled the distance between nontargets and the target in same-side versus opposite-side conditions to avoid this distance confound, and we still found this discrepancy between same-side and opposite-side conditions.

Why did saccade-related bias and NT bias *not* add up in the same-side condition? One possible explanation is that certain mechanisms exist individually or together preventing the response from getting too far away from the memorized target location. For example, other extra-retinal mechanisms for visual stability, e.g., remapping [3,4,48], might contribute to accurate target localization, and visuomotor feedback systems [49] might also contribute to accurate localization. These mechanisms might function to maintain a maximum level of error tolerance, and as a result, they might prevent the total bias from exceeding that threshold. This possibility can also explain why nontargets located on the same side as the initial fixation still facilitated response performance by reducing response variability (as shown by the size of the ellipses in Fig 2), even while they did not further bias responses.

Another possibility is that the information about nontargets on the opposite side was utilized so that it counteracted saccade-related bias, but that on the same side was somehow disregarded. As discussed before, this could be because nontargets and the initial fixation location on the same side were grouped together or provided similar/redundant information. In the real world, we often have multiple nontargets which rarely appear only on the same side. We may be able to achieve accurate target localization by incorporating nontarget information from different locations, and/or by selectively utilizing nontargets in locations that can provide non-redundant information and potentially help most with localization.

### Nontarget locations in different reference frames

In Experiments 2 and 3, we presented nontargets before the saccade was triggered, and manipulated the NT locations to see whether nontargets in different reference frames would have different effects. We found that compared to the Baseline experiment, the Relative condition (same NT location relative to target) showed a similar amount of NT facilitation, while the Absolute condition (same absolute NT location on screen) showed less facilitation, in terms of both error magnitude and response variability. In addition, the nontarget bias was larger in the Relative condition; in the Relative condition, the nontarget bias overcompensated for the saccade-related bias when they were on opposite sides of the target, while in the Absolute condition, the NT bias did not even fully counteract the saccade bias. In general, for both facilitation and bias effects, the reference frame did not change the overall pattern of the results, but rather modulated the pattern seen in the Baseline condition. One interpretation aligned with previous literature is that the critical information for target localization across saccades was already present in the baseline condition: i.e., the relative spatial information between the saccade target and nontargets, at the time right after the saccade target was presented [10,33]. In the Relative condition, this relative spatial information was also preserved across saccades, likely enhancing the influence of the nontargets, whereas in the Absolute condition, this relative spatial information was not maintained, possibly reducing the influence of the nontargets.

The importance of relative spatial information that we found is consistent with Deubel’s finding on the effect of nontarget/landmark displacement [10]. In their study, a displacement of the landmarks broke the relative spatial information between landmarks and the target. Under the assumption that the landmarks are typically stable and unchanged, participants therefore tended to report the target to be displaced in the opposite direction. Our results provide converging evidence that the relative spatial information between nontargets and the target is important, not only to decide whether the target was displaced or not, but also to recall the specific target location. While it may seem somewhat counterintuitive that landmarks are more influential when they move with the eyes to preserve relative position, rather than remain stable in environmental or absolute coordinates, this idea is also consistent with a related retinotopic benefit phenomenon, such as spatial attention lingering in retinotopic coordinates after a saccade [35], and more precise memory for retinotopic than spatiotopic locations [36,47]. Note that in our study, the peripheral nontargets in the “relative” condition were not strictly retinotopic, since they moved with the saccade target cue rather than the actual eye position. Thus, during the saccade, the retinotopic locations of the NTs were constantly changing, but the critical *relative* spatial location between the target and NTs was maintained.

It should be noted that there was a confound in the Absolute experiment that could potentially lead to a weaker NT effect than the other two experiments. As described above, we attempted to control the distance between the nontargets and the target when the initial fixation location and NT location were on the same side versus opposite sides. However, the only way this was possible in the Absolute condition was to vary the initial nontarget-target distance, resulting in an overall greater average distance for Absolute trials. Previous studies have demonstrated that larger distances between nontargets and the target could reduce the influence of nontargets on target localization [10]. Thus, it is possible that the larger average distance in Absolute experiment contributed to the weaker effects. However, even when we looked at trials in which the NT-target distance was restricted to the equivalent “near-near” cases only, there was still greater facilitation for Relative than Absolute conditions, a result indicating an effect of reference frame on top of the distance effect. Moreover, it is worth emphasizing that the existence of a distance effect itself is another example of the importance of *relative* distance to the target.

### Landmarks or distractors?

As discussed above, our results showed that the presence of nontargets both decreased response variability and induced response bias. Did the presence of nontargets actually help with or hurt target localization? In our study, overall nontargets facilitated performance; on average the responses were closer to the correct location when nontargets were presented, suggesting that the nontargets served as helpful landmarks. But it is also possible that the nontargets acted as distractors, because the responses were biased with smaller variability, as if participants responded more consistently at a wrong location. A related open question is whether subjects were consciously using the nontargets as landmarks to have a more accurate location in mind, and further, whether the presence of nontargets influenced where participants were perceiving the target to be (perceptual bias), and/or where they were clicking the mouse during the decision phase (response bias).

Future studies may investigate more into the above two interpretations, to further our understanding of the internal representation of target location. In addition, future work may manipulate the physical properties (e.g., similarity, salience, location, validity) of multiple independent nontargets, to explore how various types of NT information can be incorporated in different real-world scenarios.

## Conclusion

In summary, our experiments showed that the presence of nontargets influenced target localization. This influence seemed to manifest as a general effect on target localization rather than something specific to saccade-related processing. We argue that during a localization task - with or without saccade - the spatial location of the target is memorized along with the relative spatial information between the target and nontargets. This information may be stored in memory to reduce response variability, but the information can be distorted such that it induces a response bias at the same time. If the target localization is done across a saccade, the saccade trajectory (initial fixation location and current eye position) might also be stored as spatial references to potentially benefit and/or bias responses, and pre-saccadic and post-saccadic memories are likely incorporated together. Our representation of the target location is thus influenced by a combination of these factors - perhaps weighed by the most non-redundant information - to produce behavioral responses.

## Acknowledgement

The authors thank Emma Wu Dowd for inspiring discussion during writing.

## Supporting information

**S1 Fig. Influence of saccade landing position on (saccade-related) response bias.** A) Saccade landing position. Data are shown for saccade trials with 0 nontargets. Positive values indicate saccade landing positions biased towards the initial fixation location (i.e., undershoot). There was a significant saccade undershoot on average when there was zero NT, t(47)=11.33, p<.001, Cohen’s d=l.635. B) Saccade-related response bias. Here bias (directional errors) is shown for no-saccade and saccade trials with 0 nontargets. Positive values indicate a response bias towards the initial fixation location on saccade trials, and towards right on no-saccade trials. Data here are replotted from main test Figure 4, 0-NT. C) Saccade-related response bias in undershoot trials and overshoot trials. The saccade trials in (B) were separated into undershoot trials and overshoot trials based on saccade landing position. Again, positive values indicate a response bias towards the initial fixation location, and this bias was found in both undershot and overshot trials, with only a difference in the magnitude of the bias. The schematic above shows the scenarios indicated by the results. Arrows show the direction of saccades; eye symbols indicate the saccade landing positions; red crosses indicate the correct target locations; black crosses show the actual response locations. This part of data was submitted to 2 (saccade landing position: undershoot, overshoot) × 3 (experiment: 1, 2, 3) mixed-design ANOVA. There was a significant main effect of saccade landing position, F(1,45)=9.102, p=.004, η_p_^2^=.168, but no interaction between saccade landing position and experiment, F(2,45)=0.035, p=.965, η_p_^2^=.002. The response bias was indeed smaller on overshoot trials compared to undershoot trials, but it was still significantly greater than zero in each experiment, t’s≥2.802, p’s≤.013, Cohen’s d’s≥0.700 (p values corrected for multiple comparisons). N=16 for each experiment. Error bars are SEM.

**S2 Fig. Influence of nontarget number on eye position**. A) Eye position error magnitude defined as the distance between the final eye position just before response period and the correct fixation/saccade target location, incorporating error on both horizontal and vertical axes. Compare to main text Figure 3A (influence of nontarget number on manual target localization response accuracy). Whereas nontargets decreased manual target localization error (improving performance), the same pattern was not found for eye position (oculomotor) accuracy. A 2 (saccade presence: 0, 1) × 3 (NT number: 0, 1, 2) × 3 (experiment: 1, 2, 3) mixed-design ANOVA found a significant main effect of NT number, that the presence of NT(s) actually increased eye position error, F(2,90)=11.892, p<.001, η_p_^2^=.209. B) Similar to A) but on eye position variability, calculated using RMSD. Compared to main text Figure 3B where nontargets decreased manual response variability, nontargets significantly increased eye position variability, F(1.357,61.077)=3.690, p=.047, η_p_^2^=.076. N=16 for each experiment. Error bars are SEM.

**S3 Fig. Different distance conditions in Absolute experiment compared to Baseline and Relative experiments.** A) Schematic showing same-side and opposite-side conditions for Relative and Absolute experiments (example here shows rightward saccades). Black cross and black circles indicate initial fixation location and initial NT positions; white cross and solid circles indicate final fixation location and final NT positions. B) Descriptive scatter plots show the response distribution and the 95% confidence ellipse, as in main text Figure 2, but here plotted separate for each distance condition in Absolute experiment. Data are collapsed across participants for visualization; N=16 for each experiment.

**S4 Fig. Influence of different distance conditions on response error magnitude and variability, separating the same-and opposite-side conditions.** A) Error magnitude comparisons between three distance conditions in Absolute experiment, as well as conditions in the Baseline and Relative experiments. Left figure shows opposite-side conditions, and right figure shows same-side conditions. As in the main text, we calculated “NT facilitation” as the difference in response error magnitude for 1 and 2 NTs compared to the zero NT trials in the same experiment/condition. For the Absolute same-side conditions, a 2 (distance: near-far, near-near) × 2 (NT number: 1, 2) mixed-design ANOVA on these facilitation scores reported no significant main effects of distance or NT number, nor interaction, F’s≤2.069, p’s≥.171, η_p_^2^’s≤0.121. There was also no significant difference between absolute-same near-near and relative-same (also near-near), F(1,15)=2.621, p=.126, η_p_^2^=.149. B) Similar analyses to A) but for response variability. Here there was a significant main effect of distance, F(1,15)=5.432, p=.034, η_p_^2^=.266, with stronger facilitation for Abs-same near-near than Abs-same near-far. There was also a significant difference between Abs-same near-near and relative-same (near-near), F(1,15)=10.978, p=.005, η_p_^2^=.423, revealing an effect of reference frame on top of the distance effect. N=16 for each experiment. Error bars are SEM.

